# High-throughput single-cell chromatin accessibility CRISPR screens enable unbiased identification of regulatory networks in cancer

**DOI:** 10.1101/2020.11.02.364265

**Authors:** Sarah E. Pierce, Jeffrey M. Granja, William J. Greenleaf

## Abstract

Spear-ATAC is a modified droplet-based single-cell ATAC-seq (scATAC-seq) protocol that enables simultaneous read-out of chromatin accessibility profiles and integrated sgRNA spacer sequences from thousands of individual cells at a time. Spear-ATAC profiling of 104,592 cells representing 414 sgRNA knock-down populations revealed the temporal dynamics of epigenetic responses to regulatory perturbations in cancer cells and the associations between transcription factor binding profiles, demonstrating a high-throughput method for perturbing and evaluating dynamic single-cell epigenetic states.

## Main

Complex epigenetic regulation is a unique requirement for all multicellular organisms, enabling diverse phenotypes stemming from the same underlying genotype. Understanding how transcription factor binding dynamics drive epigenetic states is essential for the molecular dissection of many core processes, including embryogenesis, differentiation, and cancer^1^. Perturbing the expression levels of epigenetic regulators and observing the subsequent effects on chromatin accessibility provides a powerful means to study transcription factor function. To this end, CRISPR/Cas9 technologies enable precise tuning of gene expression levels using targeted mutation and epigenetic modulation strategies^2,3^. Combined with technologies such as scRNA-seq, the global transcriptional effects of these perturbations can be assayed across thousands of cells at once^4–6^. In contrast, however, current methods to profile the effects of CRISPR/Cas9-based perturbations on single-cell epigenomes are limited to analyzing 96 cells per run on an integrated fluidic circuit^7^. Here, we developed the droplet-based Spear-ATAC protocol (<u>S</u>ingle-cell <u>pe</u>rturbations with an <u>a</u>ccessibility <u>r</u>ead-out using sc<u>ATAC</u>-seq) to quantify and map the effects of perturbing transcription factor levels on chromatin accessibility in high throughput. In contrast to previous methods, Spear-ATAC relies on reading out sgRNA spacer sequences directly from genomic DNA rather than off of RNA transcripts.

Similar to bulk accessibility profiling using ATAC-seq^8,9^, the droplet-based scATAC-seq protocol begins with nuclei isolation and transposition of the sample of interest using a hyperactive transposase (Tn5) that integrates into areas of open chromatin^10,11^ (**Fig. 1a** and **Extended Data Fig. 1a**). To first ensure that capture of a single-copy of an integrated sgRNA does not depend on the local accessibility context surrounding the sgRNA sequence, we flanked the lentiviral sgRNA spacer with pre-integrated Nextera Read1 and Read2 adapters (**Extended Data Fig. 2a-b**), enabling the sgRNA sequence to be amplified with the same primers used to amplify ATAC-seq fragments in the library. When testing the detection of sgRNA fragments with bulk ATAC-seq, this design increased our ability to detect sgRNA fragments by ~4-fold without altering sgRNA efficacy (**Extended Data Fig. 2c-d**). Following transposition, the nuclei are loaded into the 10x Controller for the capture of individual nuclei into nanoliter-scale gel-beads in emulsion (GEMs). These GEMs contain barcoded Forward oligos complementary to the Nextera Read1 adapter to amplify all ATAC fragments, thereby tagging each ATAC fragment from the same nucleus with the same 10x barcode. Since this protocol uses a single barcoded primer to tag ATAC fragments, we reasoned that we could preferentially amplify each sgRNA fragment by also spiking in a Reverse oligo specific to the sgRNA backbone (**Fig 1a** and **Extended Data Fig. 3a-b**). This modification allows for exponential amplification of the sgRNA fragment at the same time the rest of the library is amplified linearly, while still ensuring the sgRNA can be used as a substrate for the second round of PCR. We also extended the number of cycles of in-GEM linear amplification of scATAC-seq fragments from 12 to 15, which subsequently adds three rounds of exponential sgRNA amplification without altering scATAC-seq quality (**Extended Data Fig. 3c-d**). Finally, we included a biotin-tagged primer during targeted sgRNA amplification which allows for the specific enrichment of these fragments while minimizing aberrant background signal from scATAC-seq reads (**Extended Data Fig. 3e**). Overall, these changes increase our ability to detect sgRNA fragments by ~40-fold compared to lentiviral integration alone followed by traditional droplet-based scATAC-seq.

**Figure 1.**
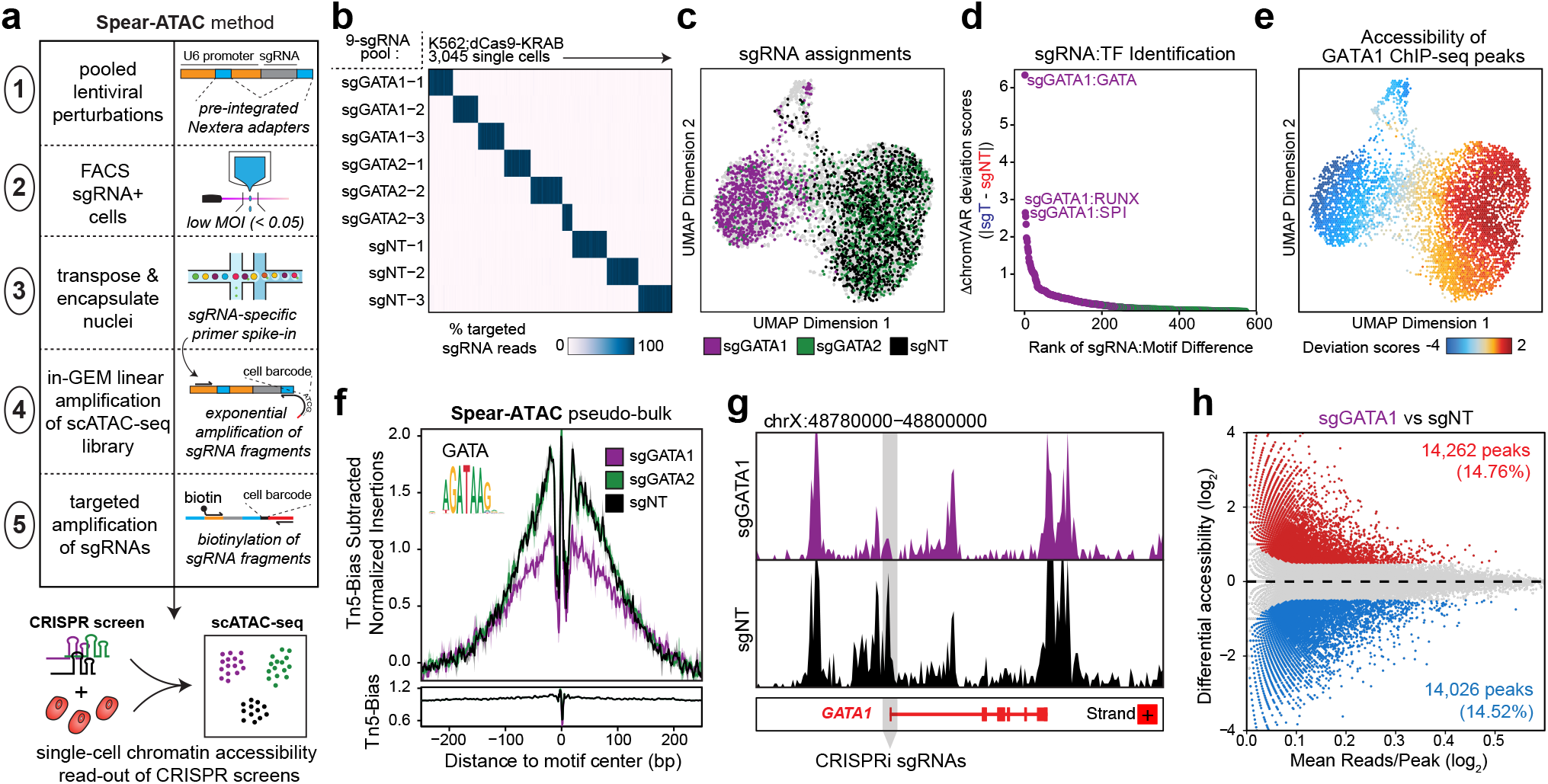
Spear-ATAC enables high-throughput CRISPR screening with a chromatin accessibility read-out. **a.** Schematic of the Spear-ATAC method. Modifications to traditional CRISPR screening methods and scATAC-seq approaches are outlined on the right. **b.** Heatmap of the percent of sgRNA reads assigned to 3,045 individual cells with corresponding chromatin accessibility information via scATAC-seq. **c.** UMAP of Spear-ATAC chromatin accessibility profiles for the pilot K562 screen colored by sgRNA assignments. Cells unassigned are in light grey. **d.** Rank ordered plot of sgRNA:TF perturbations to identify top hits in the pilot K562 screen. **e.** UMAP of Spear-ATAC chromatin accessibility profiles for the pilot K562 screen colored by chromVAR deviations for GATA1 ENCODE ChIP-seq. **f.** (*Top*) Bias-Normalized footprint of the local accessibility for each scATAC-seq cluster for genomic regions containing GATA motifs. (*Bottom*) Modeled hexamer insertion bias of Tn5 around sites containing each motif. **g.** Pseudo-bulk ATAC-seq track at the GATA1 locus for sgGATA1 and sgNT cells. Light grey box indicates the region targeted by sgGATA1-1, sgGATA1-2, and sgGATA1-2 CRISPRi sgRNAs. **h.** Differential accessibility between sgGATA1 and sgNT cells. The x-axis represents the log2 mean accessibility per peak and the y-axis represents the log2 fold change in sgGATA1 cells compared to sgNT cells. Colored points are significant (|log_2_ fold change|>0.5, FDR <0.05).

We first piloted the Spear-ATAC method with a pool of nine CRISPRi sgRNAs targeting two transcription factors (GATA1 and GATA2) and three inert sgRNA controls (Non-targeting or NT) (**Extended Data Fig. 4a**). We introduced this library into K562 leukemia cells engineered to express a CRISPRi dCas9-KRAB cassette to knockdown genes of interest, expanded the cells for six days, and then FACS-isolated sgRNA^+^ cells to process for Spear-ATAC. We captured 6,390 nuclei in the pilot run, of which we were able to directly associate 48% of single-cell epigenetic profiles (n=3,045 nuclei) to their appropriate sgRNA target with >80% specificity (**Fig 1b** and **Extended Data Fig. 4b-e**). Capturing the same number of cell-sgRNA assignments with existing methods would have required ~30 Perturb-ATAC runs costing $9.80/cell compared to one Spear-ATAC run costing $0.46/cell. Perturb-ATAC also requires 4-hour run times on a Fluidigm C1 to process each set of 96 cells, necessitating the handling of multiple batches of frozen cells over several days. Apart from the use of standard PCR machines and bead purification steps, Spear-ATAC only requires an 8-minute run time on a 10x Controller to process up to 80,000 nuclei at once, greatly increasing the potential throughput of these methods.

From the 3,045 nuclei assigned to sgRNAs in the pilot Spear-ATAC run, Uniform Manifold Approximation and Projection (UMAP) clearly distinguished cells harboring sgGATA1 from both sgGATA2 and sgNT cells, indicating the high specificity of sgRNA assignments (**Fig. 1c** and **Extended Data Fig. 4c**). GATA1 and GATA2 are both involved in hematopoietic differentiation and development; however, the erythroid transcription factor GATA1 is specifically an essential gene in K562 cells, whereas GATA2 is dispensable for growth and survival in this cell line^12^. We next developed a framework for unbiased identification of changes in TF motif accessibility across multiple populations of cells harboring different sgRNAs. We first computed TF motif accessibility scores (e.g. chromVAR deviation scores^13^ across all single cells for a given sgRNA genotype (sgT), then subtracted the average TF motif accessibility scores of the non-targeting (sgNT) cells. We then ranked all of these sgRNA-to-TF motif accessibility difference scores (sgRNA:TF scores) to identify hits. As would be expected, knockdown of GATA1 decreased the accessibility of peaks containing the GATA motif, as well as the accessibility of peaks overlapping with known GATA1 ChIP-seq peaks. Furthermore, GATA1 knockdown resulted in a muted GATA footprint compared to K562;dCas9-KRAB cells expressing non-targeting sgRNAs (**Fig. 1d-f**). Local accessibility at the GATA1 locus also decreased following knockdown, further validating that cells assigned to sgGATA1 are down-regulating expression at this locus (**Fig. 1g** and **Extended Data Fig. 5a**). Interestingly, knocking down GATA1 led to a modest increase in accessibility of GATA3 ChIP-seq peaks as well as an increase in local accessibility near the GATA3 locus (**Extended Data Fig. 5b-c**). GATA3 is typically active in the lymphoid lineage^14^ and is not expressed in K562 cells at baseline, suggesting that GATA3 is specifically activated in response to GATA1 knock down.

By performing differential accessibility analysis between sgGATA1-containing cells and sgNT-expressing control cells, we observed 14,262 peaks (14.76%) increasing in accessibility and 14,026 peaks (14.52%) decreasing in accessibility (**Fig. 1h**). Each of the three sgRNAs targeting GATA1 resulted in nearly indistinguishable chromatin accessibility profiles, with peaks decreasing in accessibility following GATA1 knockdown enriched for the GATA motif and peaks increasing in accessibility following GATA1 knockdown enriched for SPI/RUNX motifs (**Extended Data Fig. 5d**). Furthermore, individual cells with the lowest aggregate accessibility of genomic regions containing GATA1 motifs had the highest aggregate accessibility of genomic regions containing SPI/RUNX motifs and vice versa, further underscoring these regulatory relationships (**Extended Data Fig. 5e-f**). Supporting these observations, SPI (also known as PU.1) and GATA1 have been previously shown to physically interact and negatively regulate each other^15^, exemplifying the type of direct phenotypes that can be assayed and validated using the Spear-ATAC method.

Beyond motif enrichment, GREAT^16^ enrichment of genomic regions which decreased in accessibility following GATA1 knockdown were enriched for being near erythroid-specific genes (**Extended Data Fig. 5g**), and genomic regions which increased in accessibility following GATA1 knockdown were enriched for being near megakaryocyte-specific genes (**Extended Data Fig. 5h**). This result is particularly interesting given that K562 cells are often used as a model system for erythro-megakaryocytic progenitor cells. Therefore, knocking down GATA1 in K562 leukemia cells appears to prematurely push cells down a more SP1/RUNX1+, megakaryocyte lineage, despite the fact that GATA1 activity is typically required for this differentiation process^14^. Consistent with this idea, genetic disorders that impair GATA1 function often result in both the dysregulation of erythropoiesis as well as an increased incidence of transient myeloproliferative disorder and/or acute megakaryoblastic leukemia in a subset of patients^17^.

We next took advantage of the throughput of Spear-ATAC to map the dynamic effects of knocking down transcription factors over time. Traditional proliferation based CRISPR screens evaluate the representation of sgRNAs after up to three weeks in culture; therefore, we evaluated knockdown profiles 3, 6, 9, and 21 days post-knockdown. We introduced a library of 18 sgRNAs targeting 6 transcription factors (n=3 sgRNAs each) as well as 3 inert sgRNA controls into K562;dCas9-KRAB cells and performed scATAC-seq across the four time-points (**Fig. 2a** and **Extended Data Fig. 6a-h**). Similar to a proliferation-based CRISPR screen, representation of sgRNAs can be monitored over time using Spear-ATAC; for example, we observed a significant reduction in the representation of sgGATA1-containing cells at days 9 and 21 compared to days 3 and 6, whereas representation of cells with guides targeting KLF1 remained constant across days 3, 6, and 9 before decreasing at day 21 (**Fig. 2b** and **Extended Data Fig. 6f**). To identify hits from the screen – i.e. guides with significant effects on the chromatin landscape – we again used chromVAR to rank TF motif accessibility changes following sgRNA perturbations (**Fig. 2c**). Motif regulatory changes following GATA1 and KLF1 knockdown were the most significant across the genotypes, although the responses to sgGATA1 diminished over time corresponding to a decrease in representation of sgGATA1 cells in the population (**Fig. 2c**). The peaks changing in accessibility also changed over time (**Fig. 2d** and **Extended Data Fig. 6j**); for example, peaks enriched for STAT5 motifs increased in accessibility soon after GATA1 knockdown at day 3 but returned to near baseline levels of accessibility at days 6 and 9 (**Fig. 2d**). STAT5 is known to be involved in the maintenance of erythroid differentiation in a GATA1-dependent process^18^; therefore, decreased accessibility of peaks containing STAT5 motifs followed by the increased accessibility of peaks containing SPI motifs at days 6 and 9 might further suggest a transition to a more megakaryocyte lineage. Local accessibility near erythroid and megakaryocytic genes also changed as a function of time following knockdown, further emphasizing the importance of timing when evaluating the effects of perturbations on chromatin accessibility (**Fig. 2e** and **Extended Data Fig. 6i**).

**Figure 2.**
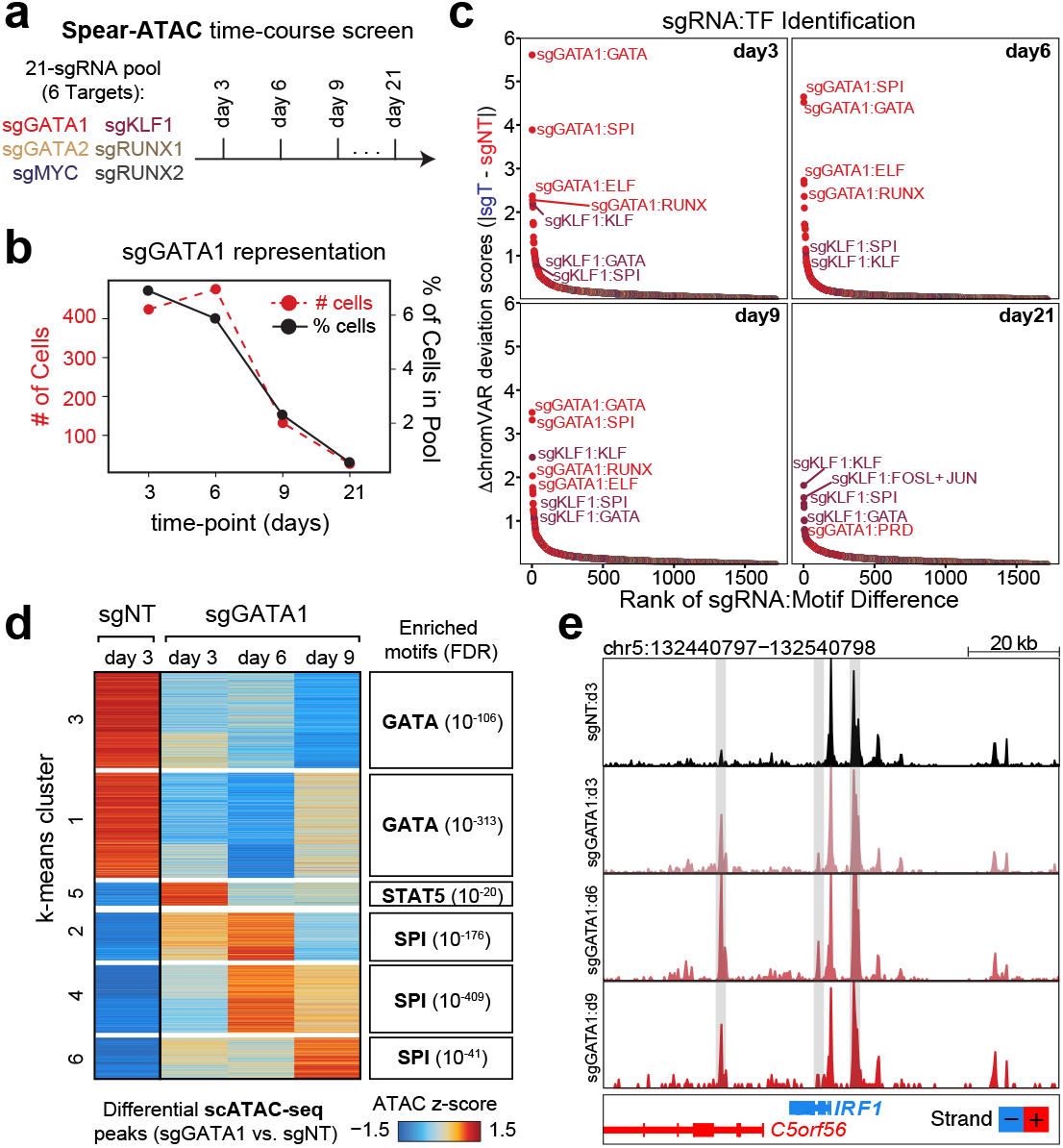
Assessing trans regulatory perturbations over time using Spear-ATAC. **a.** Schematic of Spear-ATAC time-course experiment with a 21-sgRNA pool analyzed at time-points (3 days, 6 days, 9 days, and 21 days post-transduction). **b.** Change in sgGATA1 representation over time, represented by the number of cells analyzed per time-point (red) and the % of cells in the total pool (black). **c.** Rank ordered plot of sgRNA:TF perturbations to identify top hits in the K562 time-course screen at the indicated time points. **d.** (*Left*) Heatmap of chromatin peak accessibility for each scATAC-seq sub-population using the top differential scATAC-seq peaks for sgGATA1. Each row represents a z-score of log_2_ normalized accessibility within each group using scATAC-seq. (*Right*) Transcription factor hypergeometric motif enrichment with FDR indicated in parentheses. **e.** Pseudo-bulk ATAC-seq track at the IRF1 locus for sgGATA1 (day3, day6, day9, and day21) and sgNT cells (day3). Light grey box indicates peak regions that increased in accessibility in the sgGATA1 vs sgNT cells.

Finally, to test the ability of Spear-ATAC to screen the chromatin accessibility effect of transcription factors in high-throughput, we evaluated the effects of knocking down 38 transcription factors expressed in K562;dCas9-KRAB leukemia cells with 2-3 sgRNAs each, in addition to 15 control non-targeting sgRNAs and 16 sgRNAs targeting essential genes (**Fig. 3a** and **Extended Data Fig. 7a-d**). We chose a variety of transcription factors with growth effects when knocked down in K562 cells (Growth TFs) as well as ones with no proliferation phenotype following knockdown (Non-Growth TFs)^19^ (**Fig. 3a**). Overall, we captured 32,832 nuclei representing 128 sgRNA genotypes across six Spear-ATAC samples, with on average 372 single cells being assigned to each sgRNA target with high specificity (**Extended Data Fig. 7a**). We next used chromVAR to rank motif accessibility changes following sgRNA perturbations and identified the top sgRNA:TF motif associations. We consistently identified the sgGATA1:GATA and sgKLF:KLF pairs as well as additional pairs such as sgNFE2:NFE2 and sgFOSL1:FOSL (**Fig. 3b)**. Similarly, for sgRNA genotypes that resulted in strong motif accessibility differences compared to sgNT-containing cells, the motifs identified were often consistent with the targeted transcription factor, as shown for GATA1, NFE2, KLF1, FOSL1, and NRF1 (**Fig. 3c**).

**Figure 3.**
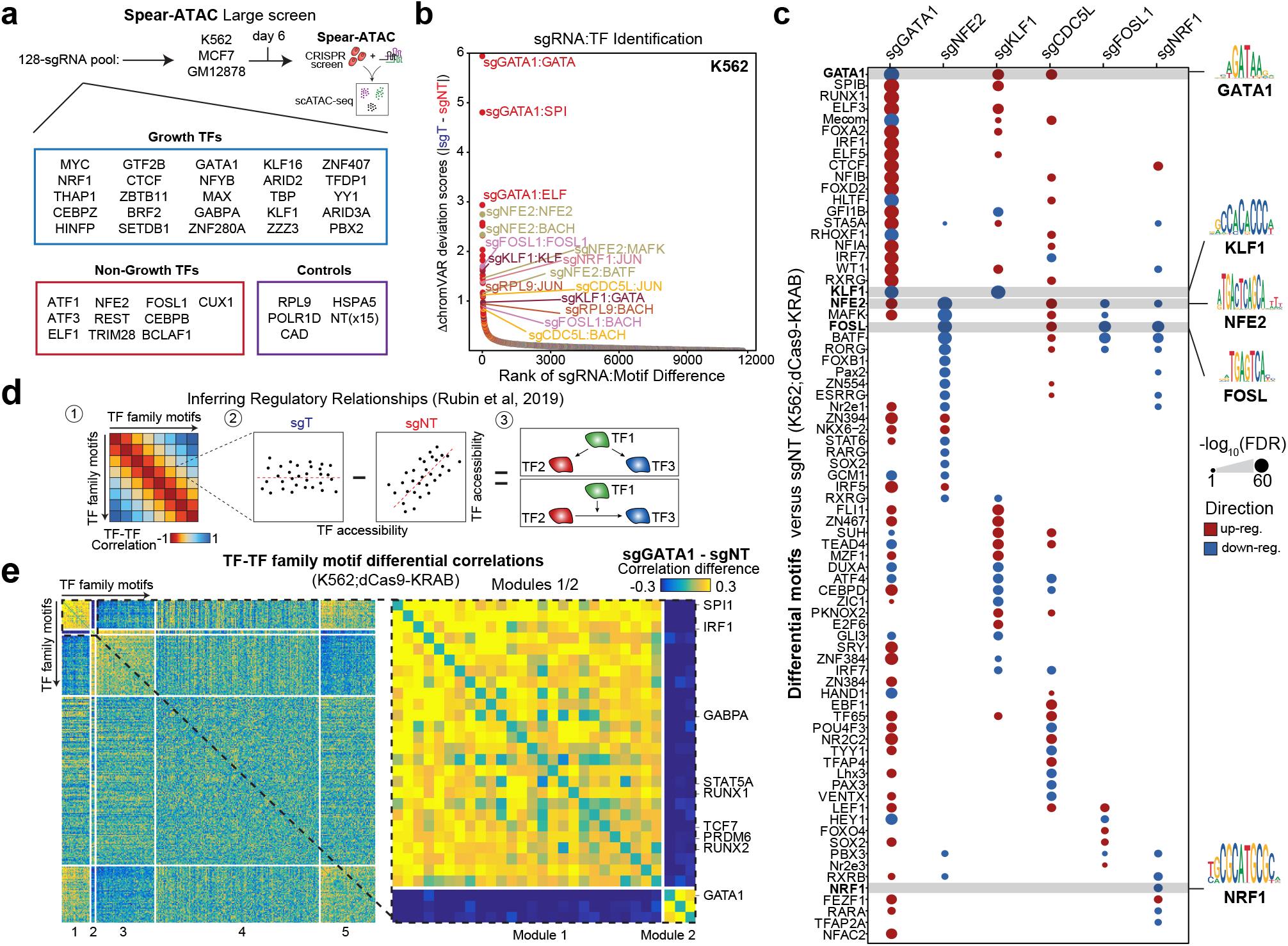
High throughput chromatin accessibility screening of CRISPR perturbations using Spear-ATAC. **a.** Schematic of Spear-ATAC large screens with a 128-sgRNA pool analyzed for 3 different cell lines (K562, MCF7 and GM12878). **b.** Rank ordered plot of sgRNA:TF perturbations to identify top hits in the K562 large screen. **c.** Motif accessibility differences across 6 perturbed transcription factors in the K562 large screen. Color indicates whether the motif accessibility difference corresponds to up-regulated (red) or down-regulated (blue) motifs. **d.** Schematic of inferring regulatory relations with Spear-ATAC. Briefly, the correlation for each TF-TF motif is determined in both the targeting (sgT) and non-targeting (sgNT) cells. Next the correlation in the non-targeting cells is subtracted from targeting cells. These TF-TF motif pairs are then assessed for different regulatory relationships. **e.** (*Left*) Heatmap of the differences between sgGATA1 targeting and non-targeting cells for all TF-TF motif accessibility correlations grouped into 5 different modules in the K562 large screen. (*Right*) Zoomed-in heatmap of modules 1 and 2 highlighting transcription factors with largely perturbed TF-TF motif accessibility relationships.

To establish regulatory relationships between TFs, we measured the effects of TF perturbations on co-varying regulatory networks^7^ (**Fig. 3d**). To identify these perturbed co-varying networks, we subtracted the TF-TF motif accessibility correlations within the non-targeting cells by the targeting cells. We first analyzed these relationships for the strongest target perturbation, sgGATA1. We identified 5 modules of TF motifs that are differentially perturbed by GATA1 knockdown (**Fig. 3e**). From this analysis, we more unbiasedly found that depletion of GATA1 led to increased coordinated activity of Module 1 consisting of crucial hematopoietic TFs such as SPI1, IRF1, RUNX and others. To further test the specificity and performance of Spear-ATAC in additional cell lines, we performed this same K562-optimized TF screen in GM12878;dCas9-KRAB lymphoblastic cells and MCF7;dCas9-KRAB breast cancer cells. Overall, we captured an additional 12,175 cells with sgRNA associations between the two cell lines. As expected, the sgRNA:TF motif perturbations were strongest and specific to K562 cells, highlighting cell-type specificity for TF regulation. However, shared patterns of co-varying regulatory networks were also uniquely observed in MCF7;dCas9-KRAB cells following knockdown of HINFP, CUX1, and NRF1 (**Extended Data Fig. 9a-c**). While HINFP, CUX1, and NRF1 have not previously been shown to directly interact with each other, HINFP and CUX1 are both involved with histone H4 gene regulation and their overlapping regulatory networks suggest a common pathway^20^.

In conclusion, Spear-ATAC can be used to evaluate the effects of perturbing transcription factor expression on gene regulatory networks, increasing the throughput of previous methods by between 35- and 100-fold (depending on target nuclei capture rate) and decreasing cost by 20-fold. An exciting application for Spear-ATAC will involve the creation of transcription factor interaction maps following multiplexed perturbations, which will enable a higher-level understanding of how proteins interact to regulate the non-coding genome. We additionally envision the application of this method following the perturbation of individual regulatory elements through high-fidelity editing methods such as prime editing^21^, allowing a quantitative understanding of how disease-related mutations alter transcription factor occupancy (as inferred by ATAC-seq) and accessibility at these sites. Spear-ATAC also enables facile monitoring of pooled epigenetic perturbations across time, providing insight into the timescales involved in epigenetic reprogramming. Given the time-dependent differences we observe in our chromatin accessibility profiles following CRISPRi perturbations, we believe that this temporal dimension of monitoring is crucial for identifying the appropriate timepoint for a given study to exclude or include downstream effects. Finally, the Spear-ATAC workflow is not inherently limited to reading out CRISPR/Cas9 sgRNAs, but could be adapted to identify sample barcodes for higher throughput multiplexing or to read-out dynamic lineage tracing marks to understand the relationship between cells during differentiation or cancer evolution.

## Materials availability

Plasmids generated in this study are available from the Lead Contact without restriction.

## Data availability

We have made available all matrices (peak matrix and chromVAR) are available through AWS (See Supplementary Table 6). We also made the 10x cell ranger atac output files and all scATAC-seq matrices used in this study available through AWS. All sequencing data have been deposited in the Gene Expression Omnibus (GEO) and are awaiting accessioning.

## Code availability

All custom code used in this work is available upon request. We additionally will host a Github website that includes the main analysis code used in this study.

## Methods

### Experimental Methods

#### Cell lines

Human cell lines (K562, GM12878, and MCF7) were a gift from Michael Bassik and Howard Chang’s laboratories, who previously purchased them from ATCC. The dCas9-KRAB derivatives used have been validated and published previously^7,22,23^. K562;dCas9-KRAB and GM12878;dCas9-KRAB cells were cultured in RPMI media supplemented with 10% FBS, 1% penicillin-streptomycin-glutamate, and 0.1% amphotericin. MCF7;dCas9-KRAB cells were cultured in DMEM media supplemented with 10% FBS, 1% penicillin-streptomycin-glutamate, and 0.1% amphotericin. All cell lines tested negative for mycoplasma using the MycoAlert Mycoplasma Detection Kit (Lonza).

#### Lentivirus production

All lentiviruses were produced by co-transfecting lentiviral backbones with packaging vectors (delta8.2 and VSV-G) into 293T cells using PEI (Polysciences). The viral-containing supernatant was collected at 48- and 72-hours post-transfection, filtered through a 0.45uM filter, and combined with fresh media to transduce cells. K562 and GM12878 derivatives were transduced by spinfection at 1000g at 37 degrees C for 2 hours. MCF7 derivatives were transduced by incubating with viral-containing supernatant for up to two days prior to the first fresh media change. Cell lines were incubated with 8ug/mL polybrene (Sigma) to enhance transduction efficiency. Cells were transduced with varying amounts of virus and Spear-ATAC was only performed on cells with an MOI < 0.05 to reduce the likelihood of multiple transduction events per cell.

#### Spear-ATAC: Cloning and modifications to the sgRNA plasmid backbone

Nextera Read1 and Read2 sequencing adapters flanking the bovine U6 promoter were inserted into the pMJ114 backbone using site-directed mutagenesis and whole-vector amplification as previously described, thereby generating pSP618. sgRNA spacer sequences of interest were inserted into the pSP618 backbone individually using site-directed mutagenesis. These sequences were originally picked from the Dolcetto CRISPRi genome-wide library available on Addgene and a full list is available in Supplementary Table 7. In addition, to allow for sgRNA read-out directly from sequencing the scATAC-seq library, we also cloned in unique, 10-bp sgRNA barcode sequences immediately adjacent to the Nextera Read1 adapter by site-directed mutagenesis (sequences also available in Supplementary Table 7). However, we found that targeted sgRNA amplification followed by targeted sgRNA sequencing gives the highest quality sgRNA:nuclei associations, and so we would recommend cloning in the sgRNA spacer sequences only and using the custom sequencing primer oMCB1672 (5’-GCCACTTTTTCAAGTTGATAACGGACTAGCCTTATTTAAACTTGCTATGCTGTTTCCAGCTTAGCTCTTAAAC-3’) for Read 1 to directly sequence the sgRNA spacer sequence. sgRNA plasmids were mixed at equimolar ratios before making virus and transducing the cells of interest.

#### Spear-ATAC: Modifications to the 10x scATAC-seq protocol

sgRNA+ nuclei were prepared as previously described for the 10x Genomics scATAC-seq protocol^10^. During GEM generation, 1.2uL of 50uM oSP1735 (5’-gctacattttacatgataggcttgg-3’) was spiked into the Master Mix. Additionally, PCR1 following GEM generation was extended from 12 cycles to 15 cycles.

#### Spear-ATAC: Amplification of sgRNA fragments out of the scATAC-seq libraries

Full protocol is described in the Supplementary Note. In brief, after the final scATAC-seq libraries were prepared (~150nM final concentration in 20uL ddH_2_O), 2.5uL of the libraries were used as input for a targeted sgRNA linear amplification PCR reaction using a 5’ biotinylated, sgRNA-specific primer (oSP2053: 5’-GTGACTGGAGTTCAGACGTGTGCTCTTCCGATCTaagtatcccttggagaaccaccttg-3’) for 25 cycles. PCR product was pooled and purified using a Qiagen Minelute kit (Qiagen). Biotinylated sgRNA fragments were then enriched using Streptavidin MyOne C1 beads (ThermoFisher) and re-suspended in 40uL ddH_2_O, which was used as input for an exponential PCR amplification reaction for 15 cycles using primers corresponding to P5 (oSP1594: 5’-AATGATACGGCGACCACCGAGA-3’) and an indexed P7-containing primer (5’-CAAGCAGAAGACGGCATACGAGAT**NNNNNNNN**GTGACTGGAGTTCAGACGTGTG-3’, where **NNNNNNNN** is replaced with the index of choice).

### Analytical Methods

#### Genome and Transcriptome Annotations

All analyses were performed with the hg38 genome. We used the hg38 genome transcripts for gene annotations from “TxDb.Hsapiens.UCSC.hg38.knownGene”.

#### SpearATAC – Aligning sgRNA data

To identify the sgRNA for each single cell we first aligned each sgRNA (conventionally Read1, i.e. for 10x scATAC “R1_001.fastq.gz”) to cell barcode (conventionally Index1, i.e. for 10x scATAC “R3_001.fastq.gz”) combination to the sgRNA library and cell barcode library respectively. We first compiled the cell barcode library of all cell barcodes with up to 1 mismatch. We additionally created a dictionary of the sgRNA barcodes using “PDict” in R. With these 2 libraries, we read in the 2 fastq reads (Read1 and Index1) in 500,000 read chunks using the package “ShortRead” in R. We next matched the Index reads to the cell barcode library using “fmatch” in R. Then, we matched the Read1 reads to the sgRNA library using “chunkDictMatch” in R. We compiled the match results into a data frame and iterated through the full fastq reads. Finally, we created a cell by sgRNA matrix that encompassed the aligned sgRNA for each cell and identified cells that had a high-fidelity sgRNA assignment as having at least 20 sgRNA counts and a specificity of 0.8 to the top target.

#### SpearATAC – Aligning scATAC data

Raw sequencing data was converted to fastq format using cellranger atac mkfastq (10x Genomics, version 1.2.0). Single-cell ATAC-seq reads were aligned to the hg38 reference genome and quantified using cellranger count (10x Genomics, version 1.2.0). The current version of Cell Ranger can be accessed here: https://support.10xgenomics.com/single-cell-atac/software/downloads/latest.

#### SpearATAC – Pre-processing scATAC data

We used ArchR^24^ (version 0.9.5) for all downstream scATAC-seq analysis (https://greenleaflab.github.io/ArchR_Website/). We used the fragments files for each sample with their corresponding csv file with cell information. We then created Arrow files using “createArrowFiles” with using the barcodes from the sample 10x CSV file with “getValidBarcodes”. This step adds the accessible fragments a genome-wide 500-bp tile matrix and a gene-score matrix. We did not filter doublets because for these screens the cells will not form many discrete clusters and thus not many heterotypic doublets can be identified. We created an ArchRProjec and then filtered cells that had a TSS enrichment below 4 and less than 1,000 fragments. For QC plots, we used “plotGroups”, “plotTSSEnrichment” and “plotFragmentSizes”. We added the sgRNA assignments for each individual sgRNA and the sgRNA targets. We reduced dimensionality with “addIterativeLSI” (default parameters), added clusters with “addClusters” (default parameters), and added a UMAP with “addUMAP” (default parameters).

To improve the fidelity of our SpearATAC sgRNA assignments, we identified the highest quality assignments for each target (similar to Replogle et al. 2020^25^). To perform this analysis, we first created an individual sgRNA by tile matrix and an sgRNA Target by tile matrix. For each target, we identified the top 5,000 increasing and 5,000 decreasing peaks between the target and non-targeting that were reproducibly regulated when comparing the individual sgRNA to the non-targeting cells. We used these 10,000 differential tile regions to perform an LSI dimensionality reduction and subsequent UMAP (n_neighbors=40, min_dist=0.4, metric=“cosine”). We next computed the “PurityRatio” for each sgRNA target cell based on the proportion of nearest neighbors being targeting cells (n = 20). Cells that had a “PurityRatio” greater than 0.9 kept their assignment (greater than 95% of assigned cells met this criterion) for downstream analysis.

Following these assignments, we created a reproducible non-overlapping peak matrix with “addGroupCoverages” and “addReproduciblePeakSet” using the sgRNA targets as groups i.e. sgGATA1, sgGATA2, sgNT, and etc. We quantified the number of Tn5 insertions per peak per cell using “addPeakMatrix”. We subsequently added motif annotations using “addMotifAnnotations” with the motifs curated and clustered from Vierstra et al 2020^26^ (https://www.vierstra.org/resources/motif_clustering). We computed chromVAR deviations for each single cell with “addDeviationsMatrix”. To identify the top sgRNA:TF perturbations, we computed the average TF motif deviations for each target and subtracted the average TF motif deviations for the non-targeting cells. By ranking the top sgRNA:TF perturbations by the absolute differences we could distinguish the top hits in each SpearATAC screen. For TF footprinting of GATA we used “plotFootprints” with normalization method “subtract” which substracts the Tn5 bias from the ATAC footprint. When performing motif based analyses, we first ranked all motifs based on variability (relevant to the analysis) and the kept the highest motif for each motif cluster/family identified from Vierstra et al 2020^26^ (https://www.vierstra.org/resources/motif_clustering). This filtration step removed redundant motifs which can confound downstream analysis.

#### SpearATAC – Analyzing K562 Pilot Screen (9-sgRNA)

Following preprocessing of the SpearATAC data, we identified differential peaks for each target vs non-target cells using “getMarkerFeatures” (testMethod=”binomial”). We identified differential peaks as those with a |log2FC| greater than 0.5 and FDR less than 0.1. Differential peaks for sgGATA1 (up-regulated and down-regulated independently) were used as input to GREAT^16^ (Association=“Two nearest genes”) to identify inferred regulated biological processes (i.e. GO terms). We next computed the average accessibility per peak for each individual sgRNA using “getGroupSE” (scaleTo=10^6). To create a heatmap of differential peaks for each sgRNA of a target with sgNT (see **Extended Data Figure 5d**), we subset by all differential peaks that were |log2FC| greater than 1 and then plotted a k-means (k=4) z-score (log2-transformed) heatmap using “ArchR:::.ArchRHeatmap”. To identify motifs enriched in each k-means cluster of peaks we used “ArchR:::.computeEnrichment” with the motifmatches and all peaks as a background set. Lastly, we computed a chromVAR deviations matrix using the ENCODE ChIP seq data set within ArchR with “addArchRAnnotations” (“EncodeTFBS”) and “addDeviationsMatrix”.

#### SpearATAC – Analyzing K562 Time-Course Screen (21-sgRNA)

Following preprocessing of the SpearATAC data, we identified differential peaks for each target vs non-target cells using “getMarkerFeatures” (testMethod=”binomial”) for each time point (day3, day6, day9 and day21). For each time point, we identified differential peaks as those with a |log2FC| greater than 0.5 and FDR less than 0.1. We next computed the average accessibility per peak for each time point and individual sgRNA using “getGroupSE” (scaleTo=10^6). To create a heatmap of differential peaks for each sgRNA of a target with sgNT (see **Figure 2d** **and Extended Data Figure 6j**), we first subset by the union of all differential peaks that were |log2FC| greater than 1 for each time point. Next, we computed the average log2 fold changes for sgRNA target vs the sgNT at that time point (using the pseudobulk matrix above). We further filtered the differential peaks by those peaks that have a |log2FC| greater than 0.25 in at least 1 time point. We plotted a k-means (k=6) z-score (log2-transformed) heatmap using “ArchR:::.ArchRHeatmap”. To identify motifs enriched in each k-means cluster of peaks we used “ArchR:::.computeEnrichment” with the motifmatches and all peaks as a background set.

#### SpearATAC – Analyzing Large Screens for K562, GM12878 and MCF7 (128-sgRNA)

Following preprocessing of the SpearATAC data, we identified differential motifs for each target vs non-target cells using “getMarkerFeatures” (testMethod=”wilcoxon”, bufferRatio=0.95, maxCells=250, useSeqnames=”z”). We filtered sgNT-5,6,8,11,12 cells prior to this differential comparison after identifying these sgRNA as outliers while performing pseudobulk PCA analysis. To identify perturbed co-varying regulatory networks (see **Figure 3e** **and Extended Data Figure 9a-c**), we first got the motif deviations matrix for each screen (K562, GM12878 and MCF7) and filtered cells corresponding to sgNT-5,6,8,11,12. Next, we computed the average motif deviation scores for each sgRNA target. We subtracted the average motif deviation scores from the non-targeting cells. We rank ordered the motifs by the maximum observed average motif deviation score difference (absolute difference) across all targets in each screen. Finally, we rank ordered the motifs by the average of these maxima across all three screens (K562, GM12878 and MCF7). We de-duplicated the motifs in each cluster (see Vierstra et al 2020^26^; https://www.vierstra.org/resources/motif_clustering) to remove redundant motifs. For each sgRNA we computed all TF-TF deviation score correlations for the non-redundant motifs. Each targeting sgRNA was then subtracted by the non-targeting sgRNA TF-TF correlations. This differential correlation matrix was subsequently hierarchal clustered with “hclust” and split into 5 modules with “cutree”. A heatmap of the differential correlations for the sgRNA targeting cells was then constructed across all modules.

## Acknowledgments

We thank M. Bassik and members of the Greenleaf and Chang laboratory for helpful comments. We thank the Stanford Shared FACS facility for technical support and A. Orantes for administrative support.

## Funding

S.E.P was supported by the NSF Graduate Research Fellowship Award and the Tobacco-Related Diseases Research Program Predoctoral Fellowship Award. This work was supported by NIH RM1-HG007735, UM1-HG009442, UM1-HG009436, and U19-AI057266 (to W.J.G.). WJG acknowledges funding from Emmerson Collective.

## Author contributions

S.E.P., J.M.G., and W.J.G conceived the project and designed the experiments. S.E.P. led the method development and experimental data production. J.M.G led the data analysis. J.M.G. and S.E.P. performed the ATAC-seq and scATAC-seq analysis. S.E.P. and J.M.G were supervised by W.J.G. and S.E.P., J.M.G., and W.J.G wrote the manuscript.

## Competing interests

W.J.G. is a consultant for 10x Genomics who has licensed IP associated with ATAC-seq. W.J.G. has additional affiliations with Guardant Health (consultant) and Protillion Biosciences (co-founder and consultant).

## Extended Data Figure Legends

**Extended Data Figure 1. Outline of Spear-ATAC protocol.**

**a.** Cells are transduced with a lentiviral sgRNA flanked by Read1/Read2 Nextera adapters, FACS sorted to exclude sgRNA-negative cells, and processed for scATAC-seq using a modified 10x Genomics droplet-based protocol. In brief, nuclei are isolated, transposed, and gel bead emulsions (GEMs) are made with individual nuclei re-suspended in Nuclei Resuspension Buffer combined with an enzymatic master mix containing a sgRNA-specific primer. GEMs are immediately subjected to in-GEM linear amplification of scATAC-seq fragments, while sgRNA fragments are subjected to exponential amplification using the sgRNA-specific primer as a Forward primer. The number of cycles of in-GEM amplification has been extended from 12 cycles (original 10x protocol) to 15 cycles (Spear-ATAC protocol). Following a series of clean-ups and exponential amplification to attach flow-cell adapters (P7) and sample indices to the ATAC-seq fragments, sgRNA fragments are specifically enriched using a biotin-conjugated oligo specific to the sgRNA fragments, and then amplified and sequenced separately.

**Extended Data Figure 2. Flanking a lentiviral sgRNA spacer with Read1/Read2 adapters increases sgRNA fragment capture efficiency following ATAC-seq.**

**a.** Schematic of traditional lentiviral sgRNA.

**b.** Schematic of modified Spear-ATAC lentiviral sgRNA. sgRNA spacer sequence is flanked by Read1/Read2 Nextera adapters, obviating the need for tn5 transposase to randomly insert sequencing adapters nearby the sgRNA sequence during an ATAC-seq transposition reaction. The Read2 adapter is inserted in a flexible region of the U6 promoter to decrease the total length of the sgRNA fragment.

**c.** Log2 fold change in sgRNA fragments amplified following bulk ATAC-seq reactions performed with the indicated numbers of input cells transposed with lentiviral sgRNAs with and without Nextera adapters flanking the sgRNA region.

**d.** Percentage of GFP negative cells following lentiviral transduction of sgGFP sequences with and without Nextera adapters flanking the sgRNA region.

**Extended Data Figure 3. Spear-ATAC modifications to the 10x scATAC-seq protocol do not hurt scATAC quality.**

**a-b.** scATAC-seq quality control metrics with and without a sgRNA-specific primer spiked into the enzymatic master mix during GEM formation on the 10x Controller. Insertion profiles (left a), TSS enrichment (right a), and average fragment distribution (b) in each sample is shown.

**c-d.** scATAC-seq quality control metrics after 12, 13, 14, and 15 cycles of in-GEM linear amplification. Insertion profiles (left c), TSS enrichment (right c), and average fragment distribution (d) in each sample is shown.

**e.** Biotin tagging and selection increases the specificity of targeted sgRNA amplification, as seen by the fragment distribution of products without (left) and with (right) the first biotin tagging step.

**Extended Data Figure 4. Quality control metrics and sgRNA assignment methods for 9-sgRNA GATA pool.**

**a.** Schematic of sgRNAs in the first proof-of-principle 9-sgRNA pool.

**b.** scATAC-seq quality control metrics for the proof-of-principle 9-sgRNA GATA pool. Average TSS enrichment (left), individual TSS enrichment x log10 number of unique fragments per cell (middle), and average insertion profiles (right inset) for the 6,390 nuclei captured.

**c.** UMAP of Spear-ATAC chromatin accessibility profiles (N = 6,390) for the 9-sgRNA K562 screen colored by Seurat graph clustering in ArchR.

**d.** Specificity to a single sgRNA spacer sequence and number of sgRNA reads (log10) following targeted amplification for each cell analyzed.

**e.** UMAP of Spear-ATAC chromatin accessibility profiles for the 9-sgRNA K562 screen colored by the individual sgRNA. Cells unassigned are in light grey.

**Extended Data Figure 5. GATA1 maintains accessibility at genomic regions important for erythropoiesis and knocking down GATA1 results in altered regulation of GATA3.**

**a.** Pseudo-bulk ATAC-seq track at the GATA1 locus for individual sgRNAs of sgGATA1 and sgNT. Light grey box indicates the region targeted by sgGATA1-1, sgGATA1-2, and sgGATA1-2 CRISPRi sgRNAs.

**b.** UMAP of Spear-ATAC chromatin accessibility profiles for the pilot K562 screen colored by chromVAR deviations for GATA3 ENCODE ChIP-seq.

**c.** Pseudo-bulk ATAC-seq track at the GATA3 locus for sgGATA1 and sgNT cells. Light grey boxes indicate regions that are significantly changing in accessibility following sgGATA1 knockdown.

**d.** Heatmap of chromatin peak accessibility for each individual sgRNA for both sgGATA1 and sgNT using the top differential scATAC-seq peaks identified from ArchR. Each row represents a z-score of log2 normalized accessibility within each group using scATAC-seq. (Right) Transcription factor hypergeometric motif enrichment with FDR indicated in parentheses.

**e.** Ridge plot for chromVAR deviations (normalized to the median sgNT) of individual cells for (Left) GATA, (Middle) RUNX and (Right) SPI.

**f.** ChromVAR motif correlations for the indicated transcription factor families (RUNX, SPI, or GATA). Pearson correlation (R) is indicated in the top left corner. Each individual point represents a single cell. Cells are colored based on their sgRNA genotype, as indicated in Figure 5b.

**g.** GREAT enrichment of peaks that decrease in accessibility with sgGATA1. Enrichments are taken from the Mouse Phenotype with Single KO (Knock-out) dataset.

**h.** GREAT enrichment of peaks that increase in accessibility with sgGATA1. Enrichments are taken from the GO Process dataset.

**Extended Data Figure 6. Spear-ATAC increases the throughput of time-course screens to understand the timing of effects following transcription factor perturbations.**

**a.** Schematic of 21 sgRNAs used in the Spear-ATAC K562 time-course screen.

**b.** Specificity to a single sgRNA spacer sequence and number of sgRNA reads (log10) following targeted amplification for each cell analyzed.

**c.** TSS enrichment scores per single cell across all Spear-ATAC K562 time-course time points.

**d.** Aggregate TSS enrichment scores across all Spear-ATAC K562 time-course time points.

**e.** Aggregate ATAC-seq fragment size distributions across all Spear-ATAC K562 time-course time points.

**f.** Aggregate ATAC-seq fragment size distributions across all Spear-ATAC K562 time-course time points.

**g.** Change in sgKLF1 representation over time, represented by the number of cells analyzed per time-point (orange) and the % of cells in the total pool (black). Cells unassigned are in light grey.

**h.** Ridge plot for chromVAR deviations (normalized to the median sgNT) of individual cells across each time point and target sgRNA for (Left) GATA, (Middle) SPI and (Right) KLF.

**i.** Pseudo-bulk ATAC-seq track at the (Top Left) RUNX1, (Top Right) PRKAR2B, (Bottom Left) PPBP and (Bottom Right) MPL locus for sgGATA1 (day3, day6, day9, and day21) and sgNT cells (day3). Light grey box indicates peak regions that changed in accessibility in the sgGATA1 vs sgNT cells.

**j.** (*Left*) Heatmap of chromatin peak accessibility for each scATAC-seq sub-population using the top differential scATAC-seq peaks for sgKLF1. Each row represents a z-score of log_2_ normalized accessibility within each group using scATAC-seq. (*Right*) Transcription factor hypergeometric motif enrichment with FDR indicated in parentheses.

**Extended Data Figure 7. Quality control metrics for Spear-ATAC transcription factor screens in K562;dCas9-KRAB, GM12878;dCas9-KRAB, and MCF7;dCas9-KRAB cell lines.**

**a.** Specificity to a single sgRNA spacer sequence and number of sgRNA reads (log10) following targeted amplification for each cell analyzed for the (Left) K562, (Middle) GM12878 and (Right) MCF7 large screens.

**b.** TSS enrichment scores per single cell across the (Left) K562, (Middle) GM12878 and (Right) MCF7 large screen replicates.

**c.** Aggregate ATAC-seq fragment size distributions across the (Left) K562, (Middle) GM12878 and (Right) MCF7 large screen replicates.

**d.** UMAP of Spear-ATAC chromatin accessibility profiles for the (Left) K562, (Middle) GM12878 and (Right) MCF7 large screens. This UMAP is colored by (Top) experimental replicate and (Bottom) assigned sgRNA. Cells unassigned are in light grey.

**Extended Data Figure 8. Transcription factor motif accessibility changes following perturbations in K562;dCas9-KRAB, GM12878;dCas9-KRAB, and MCF7;dCas9-KRAB cell lines.**

**a.** Rank ordered plot of sgRNA:TF perturbations to identify top hits for the (Left) K562, (Middle) GM12878 and (Right) MCF7 large screens.

**b.** Ridge plot for chromVAR deviations (normalized to the median sgNT) of individual cells for (Left) GATA, (Middle Left) NFE2, (Middle Right) KLF and (Right) FOSL across the K562 large screen.

**Extended Data Figure 9. Patterns of differential transcription factor motif correlations in MCF7;dCas9-KRAB cells following knockdown of HINFP, CUX1, and NRF1.**

**a.** (Left) Heatmap of the differences between sgHINFP targeting and non-targeting cells for all TF-TF motif accessibility correlations grouped into 5 different modules in the MCF7 large screen. (Right) Zoomed in heatmap of modules 1 and 2 highlighting transcription factors with largely perturbed TF-TF motif accessibility relationships.

**b.** (Left) Heatmap of the differences between sgCUX1 targeting and non-targeting cells for all TF-TF motif accessibility correlations grouped into 5 different modules in the MCF7 large screen. (Right) Zoomed in heatmap of modules 1 and 2 highlighting transcription factors with largely perturbed TF-TF motif accessibility relationships.

**c.** (Left) Heatmap of the differences between sgNRF1 targeting and non-targeting cells for all TF-TF motif accessibility correlations grouped into 5 different modules in the MCF7 large screen. (Right) Zoomed in heatmap of modules 1 and 2 highlighting transcription factors with largely perturbed TF-TF motif accessibility relationships.

## Supplementary Note

**Supplementary Note 1. Protocol for targeted amplification of sgRNAs out of final scATAC-seq libraries.** This supplementary note contains a full protocol for the targeted amplification of sgRNA fragments out of the final 10x scATAC-seq libraries made with Spear-ATAC.

## Supplementary Tables

**Supplementary Table 1. Spear-ATAC experiment statistics.**

This table contains information about each Spear-ATAC dataset generated in this study including QC statistics and number of sgRNA to cell associations captured.

**Supplementary Table 2. Motif accessibility results from K562 pilot Spear-ATAC screen.**

This table contains information for the sgRNA:TF motif accessibility changes between targeting and non-targeting cells for the K562 pilot screen.

**Supplementary Table 3. Motif accessibility results from K562 time-course Spear-ATAC screen.**

This table contains information for the sgRNA:TF motif accessibility changes between targeting and non-targeting cells for the K562 time-course Spear-ATAC screen.

**Supplementary Table 4. Motif accessibility results from K562, GM12878, and MCF7 transcription factor Spear-ATAC screens.**

This table contains information for the sgRNA:TF motif accessibility changes between targeting and non-targeting cells for the K562, GM12878, and MCF7 transcription factor Spear-ATAC screens.

**Supplementary Table 5. List of Vierstra motif clustering annotations used for all motif analyses.**

This table contains the motif clustering annotations used for motif de-duplication and motif based analyses.

**Supplementary Table 6. Spear-ATAC data URLs.**

This data contains information for how to access the Spear-ATAC data for each experiment in this study.

**Supplementary Table 7. List of sgRNA spacer sequences used in this study.**

This table contains information about the sgRNA spacer sequences used in this study and their targets.

